# Hydrogen sulfide modulation of intrinsic excitability in rat hippocampal CA1 pyramidal cells

**DOI:** 10.64898/2026.11.20.768207

**Authors:** Vladimir A. Martínez-Rojas, Amayrani Perez-Barragan, Luis Rojas-Hernández, Ixchel Ramírez-Camacho, David Centurion, Emilio J. Galván

## Abstract

Although traditionally known as a toxic gas, hydrogen sulfide (H_2_S) is now recognized as a biologically active molecule with modulatory effects on many physiological targets, including the central nervous system. In this context, pyramidal cells of the hippocampus express the H_2_S-generating enzymes required for its synthesis and local release. Nonetheless, the effects of H_2_S on the passive and active membrane properties of glutamatergic neurons of the hippocampus remain incompletely characterized. To investigate these effects, we examined how application of an exogenous H_2_S donor and inhibition of endogenous H_2_S synthesis shape passive membrane properties, action potential dynamics, firing frequency, and activation of voltage-dependent currents in rat hippocampal CA1 pyramidal cells. Acute exposure to the H_2_S donor sodium hydrosulfide (NaHS, 200 µM) modified passive membrane integration and action potential dynamics and reduced firing frequency. These functional alterations were associated with decreased amplitude of voltage-dependent inward and outward currents. Western blot analysis confirmed the expression of H_2_S-producing enzymes (CBS, CSE, and 3-MST) in tissue resected from the CA1 pyramidal cell layer. In contrast to NaHS, treatment with aminooxyacetic acid (AOAA, 100 µM), a nonselective inhibitor of H_2_S synthesis, produced limited effects on passive membrane properties but increased firing frequency. Collectively, these results indicate that exogenous H_2_S reduces the intrinsic excitability of hippocampal CA1 pyramidal cells. Thes findings characterize the effect of H_2_S on membrane properties and voltage-dependent currents that contribute to neuronal firing.

**Highlights:**

- Exogenous H_2_S suppresses intrinsic excitability in rat CA1 pyramidal cells.
- H_2_S reversibly reduces voltage-gated inward and outward current amplitudes.
- CBS, CSE, and 3-MST are detected in tissue resected from CA1 stratum pyramidale.
- AOAA, a nonselective inhibitor H_2_S synthesis, increases CA1 neuronal firing.

## 1. Introduction

Hydrogen sulfide (H_2_S) has emerged as an endogenous signaling molecule with diverse physiological roles across multiple organ systems, including the central nervous system (Cirino et al., 2023). Despite its well-documented toxicity at high concentrations (Ng et al., 2019), H₂S is endogenously synthesized under physiological conditions (Zhong et al., 2020). H₂S biosynthesis is linked to sulfur amino acid metabolism through enzymatic pathways involving cystathionine β-synthase (CBS), cystathionine γ-lyase (CSE), and 3-mercaptopyruvate sulfurtransferase (3-MST) (Polhemus and Lefer, 2014).

In the brain, H_2_S exerts neuroprotective effects in experimental models of oxidative stress and neuronal injury, including reduced hippocampal neuronal apoptosis (Kimura and Kimura, 2004; Kamat et al., 2015; Wang et al., 2025). Its involvement in synaptic remodeling has also been reviewed (Kamat et al., 2015). Its actions also extend to hippocampal synaptic physiology, where H_2_S enhances N-methyl-D-Aspartate (NMDA) receptor-mediated responses and facilitates the induction of long-term potentiation (Abe and Kimura, 1996). More recently, Furuie et al. (2023) demonstrated that exogenous H_2_S and hydrogen disulfide (H_2_S_2_) promote the release of glutamate, GABA, and D-serine in the rat hippocampus. In area CA1, H_2_S has also been shown to protect neurons against lead-induced damage in rats (Rafaiee et al., 2021). Collectively, these findings support a role for H₂S in hippocampal physiology and provide a rationale for examining its effects on the intrinsic excitability of CA1 pyramidal cells (PCs), a major output population of the hippocampal circuit (van Strien et al., 2009).

The activity of hippocampal circuits is shaped not only by synaptic interactions but also by the intrinsic electrical properties of their constituent neurons. In CA1 PCs action potential (AP) firing patterns depend on both depolarizing drive and intrinsic membrane properties. This relationship partially reflects, the interplay of voltage-dependent ionic conductances that determine spike initiation, repolarization, and sustained firing (Bean, 2007). H_2_S inhibits voltage-gated K^+^ currents in cultured hippocampal neurons (Dallas et al., 2021) and alters voltage-gated Na^+^ and Ca^2+^ currents in subfornical organ neurons (Kuksis and Ferguson, 2015), demonstrating its ability to modulate ionic conductances involved in intrinsic excitability. H_2_S exposure also suppresses electrical activity in primary cortical neuronal cultures (Kim et al., 2023). However, the effects of H_2_S on intrinsic excitability in identified CA1 PCs remains incompletely characterized. To address this question, we examined the effects of the H₂S donor NaHS on intrinsic excitability and voltage-dependent membrane currents in rat CA1 PCs. We also assessed the presence of H₂S-producing enzymes in the CA1 pyramidal cell layer and investigated the effects of aminooxyacetic acid (AOAA), a nonselective inhibitor of H_2_S biosynthesis, on neuronal excitability.

## Methods

### 2.1 Experimental animals

All animal procedures were approved by the Ethics Committee for Animal Research (CICUAL) of the Center for Research and Advanced Studies of the National Polytechnic Institute (CINVESTAV; protocol No. 0090–14). Effords were made to minimize animal suffering and the number of animals used. Procedures were also conducted in compliance with the Mexican Official Standard NOM-062-ZOO-1999 and the National Institutes of Health (NIH) guidelines for the care and use of laboratory animals. Male Wistar rats, aged 4-5 weeks and weighing 100-150 g, were used in this study. Animals were group-housed (2–3 per cage) in the institutional animal facility at 23 ± 1 °C under a 12:12-h inverted light–dark cycle, provided food and water ad libitum, and regular veterinary supervision.

### 2.2 Drugs

All salts and drugs used in this study were purchased from Sigma-Aldrich Chemical Co. (St. Louis, MO, USA).

### 2.3 Exogenous H_2_S delivery by NaHS perfusion in hippocampal slices

Exogenous H_2_S was supplied by continuous bath perfusion of acute hippocampal slices with solution containing sodium hydrosulfide (NaHS). The exposure interval was selected based on preliminary time-course experiments, which showed that electrophysiological effects were consistently detectable 5 min after perfusion onset. Previous electrophysiological studies have reported rapid responses to NaHS and emphasized the time-dependent availability of H_2_S in solution (Kuksis and Ferguson, 2015; Sitdikova et al., 2014). In solution, NaHS rapidly dissociates into Na^+^ and HS^−^; HS^−^exists in a pH-dependent equilibrium with molecular H_2_S. Because H_2_S is volatile and its concentration can decline rapidly in open experimental systems (DeLeon et al., 2012), continuous perfusion was used to maintain delivery of freshly supplied NaHS-containing solution during the exposure period. Concentrations are therefore expressed as nominal NaHS concentrations rather than measured free H_2_S concentrations.

### 2.4 Acute hippocampal slice preparation

Animals were deeply anesthetized with sodium pentobarbital (50 mg/kg, i.p.) and subsequently euthanized by decapitation using a guillotine. The skull was exposed, and a caudal-to-rostral incision was made along the occipital suture. The skull plates were carefully separated, and the brain was removed following established protocols to ensure accuracy, efficiency, and consistency (Griego et al., 2022). The tissue was rapidly transferred to a slushy sucrose-based solution containing (in mM): 210 sucrose, 2.8 KCl, 2 MgSO_4_, 1.25 Na_2_HPO_4_, 25 NaHCO_3_, 1 MgCl_2_, 1 CaCl_2_, and 10 D-glucose, saturated with carbogen (95% O_2_/5% CO_2_). Transverse brain slices (385 μm thick) containing the hippocampal formation and surrounding regions were obtained using a VT1000S vibratome (Leica, Nussloch, Germany). Slices were incubated at 34°C for 30 min in a holding solution continuously bubbled with carbogen and containing (in mM): 125 NaCl, 2.5 KCl, 1.25 Na_2_HPO_4_, 26 NaHCO_3_, 4 MgCl_2_, 1 CaCl_2_, and 10 D-glucose. After a 1-h recovery period, individual slices were transferred to a submerged recording chamber (total volume: 2 mL) mounted on an upright Nikon FN1 microscope (Nikon Corporation, Minato, Tokyo, Japan). Slices were continuously perfused with standard artificial cerebrospinal fluid (aCSF; ≈3.0 mL/min) using a peristaltic pump (120S, Watson-Marlow, Wilmington, MA, USA). The aCSF contained (in mM): 125 NaCl, 2.5 KCl, 1.25 Na_2_HPO_4_, 25 NaHCO_3_, 2 MgCl_2_, 2 CaCl_2_, and 10 D-glucose. For all solutions, pH was adjusted to 7.30–7.35, and osmolarity was adjusted to 300 ± 5 mOsm/kg. Recordings were performed at 32.0 ± 1 °C using an in-line solution heater connected to a TC-324C single-channel temperature controller (Warner Instruments, Hamden, CT, USA).

### 2.5 Whole-cell patch-clamp recordings

Electrophysiological recordings were performed exclusively in dorsal hippocampal slices visualized using infrared differential interference contrast (IR-DIC) optics on an Eclipse FN1 microscope (Nikon Corporation, Minato, Tokyo, Japan). CA1 PCs were identified based on their morphology and selected from the superficial portion of the stratum pyramidale. Patch pipettes were pulled from borosilicate glass capillaries using a P-97 micropipette puller (Sutter Instruments, Novato, CA, USA). Pipette-tip resistance was 3-5 MΩ when filled with an intracellular solution containing (in mM): 135 potassium gluconate, 10 KCl, 5 NaCl, 1 EGTA, 10 HEPES, 2 Mg-ATP, 0.4 Na-GTP, and 10 phosphocreatine. The intracellular solution was adjusted to a pH of 7.2–7.3 and an osmolarity of 295 mOsm/kg. Whole-cell recordings were obtained using an Axopatch 200B amplifier (Molecular Devices, San Jose, CA, USA), and signals were low-pass filtered at 2 kHz and digitized at 10 kHz using a Digidata 1550B. Voltage and current commands were generated using pCLAMP software version 10.6 (Molecular Devices). Series resistance (≈10–20 MΩ) was monitored throughout all experiments, and recordings from cells showing changes greater than 20% were excluded from analysis. Cells were excluded if their initial resting membrane potential (RMP), measured immediately after membrane break-in to the whole-cell configuration was more depolarized than −55 mV.

To limit the contribution of individual animals and avoid repeated drug exposure, only two slices per animal were analyzed, with one neuron recorded per slice and each slice subjected to a single drug perfusion.

### 2.6 Electrophysiological assessment of intrinsic membrane properties

After establishing the whole-cell configuration, neurons were allowed to stabilize for 2–3 min. Somatic input resistance was determined from voltage responses elicited by 1-s hyperpolarizing current steps delivered from 0 to −300 pA in 30-pA decrements. The membrane time constant was estimated from the voltage response evoked by a −30-pA current injection by fitting the decay phase with a single-exponential function: *f(t) = Ae^−t/r^+ C.* Membrane capacitance was estimated as Cm = τ/*Rin*. The sag ratio was calculated as the ratio of the steady-state to the peak hyperpolarizing voltage deflection from RMP during a −300-pA current pulse.

### 2.7 AP dynamics and firing analysis

For AP analyses, neurons were maintained at −65 mV. Rheobase was estimated as the current amplitude at which the first AP occurred during a 500-ms depolarizing current ramp. AP kinetics were examined using spikes generated during trains evoked by 1-s somatic current injections of 150 pA. Mean firing frequency was calculated as the number of APs elicited during a 1-s current pulse divided by the pulse duration. Input–output (I–O) curves were generated by quantifying AP output in response to depolarizing current steps from 0 to 300 pA in 30-pA increments. The I-O relationship was fitted with a three-parameter sigmoidal function: 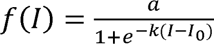 where *f(I)* stand for the firing rate as a function of the injected current *I, a* is the fitted maximal firing rate, *I_o_* the current producing half-maximal firing and *k* the steepness parameter. The offset interpreted as an extrapolated current threshold for firing initiation, was obtained from the x-intercept of a linear regression fitted to the linear portion of the I-O curve. Firing adaptation was evaluated from changes in interspike intervals (ISIs), across the spike train, with each ISI defined as the time between consecutive APs within a spike train.

### 2.8 Macroscopic current analyses

Voltage-clamp recordings were performed using a protocol adapted from Martínez-Rojas et al., (2025). Voltage-dependent membrane currents were elicited by applying a series of 500-ms voltage steps from −80 to +50 mV in 10-mV increments, from a holding potential of −65 mV. This protocol enabled the characterization of membrane currents across a range of test potentials, including transient inward and sustained outward current components. An online P/N leak-subtraction protocol was used to subtract linear leak and capacitive current components using scaled responses to small voltage pulses. The resulting currents were analyzed to assess voltage-dependent changes in membrane current.

### 2.9 Western blot

Protein expression of the H₂S-producing enzymes was examined by Western blot. Hippocampal slices from three untreated rats were used for tissue collection. Under stereomicroscopic guidance, the CA1 region was identified by the characteristic organization of its stratum pyramidale, which served as an anatomical landmark. Tissue enriched in the CA1 stratum pyramidale was isolated by manual microdissection using fine dissection scissors, minimizing the inclusion of adjacent hippocampal regions and surrounding neuropil. Microdissected tissue was homogenized in ice-cold radioimmunoprecipitation assay (RIPA) buffer containing 150 mM NaCl, 50 mM Tris (pH 8.0), 1% Triton X-100, 0.5% sodium deoxycholate, and 0.1% SDS, supplemented with an EDTA-free protease inhibitor cocktail (cOmplete™ Mini; Sigma-Aldrich). Following centrifugation to remove insoluble material, total protein concentration was determined using a Bradford protein assay. Equal amounts of protein (30 µg per sample) were resolved on 10% SDS-polyacrylamide gels at 120 V for 90 min and transferred onto polyvinylidene difluoride (PVDF) membranes at 10 V for 60 min. Membranes were blocked with 5% nonfat dry milk in phosphate-buffered saline containing 0.1% Tween 20 (PBS-T) and incubated with primary antibodies against CBS (1:500; Santa Cruz Biotechnology; Cat. No. sc-133154), CSE (1:1,000; Santa Cruz Biotechnology; Cat. No. sc-365381), or 3-MST (1:1,000; Sigma-Aldrich; Cat. No. HPA001240). β-Actin (1:10,000; GeneTex; Cat. No. GTX5512) was used as a loading control. Membranes were subsequently incubated with an anti-mouse secondary antibodies (1:7,000; GeneTex; Cat. No. GTX21311-01) for CBS, CSE, and β-actin, and an anti-rabbit secondary antibody (1:5,000; GeneTex; Cat. No. GTX213110-01) for 3-MST. Chemiluminescent signals were acquired using a ChemiDoc™ XRS+ Imaging System (Bio-Rad, CA, USA).

### 2.10 Electrophysiological data processing, statistical analysis, and data visualization

Electrophysiological recordings were processed using a custom Python-based analysis pipeline. Raw Axon Binary Format (ABF) files were imported using pyABF, and AP detection and waveform characterization were performed using the Intrinsic Physiology Feature Extractor (IPFX; Allen Institute for Brain Science). The custom Python script used for ABF processing and AP feature extraction is publicly available on GitHub (https://github.com/MiNoS-Ephys/ABF_IPFX_CA1), together with its analysis parameters and software dependencies.

Electrophysiological measurements were analyzed using a paired experimental design, with recordings obtained from the same cells before (Ctrl) and after NaHS exposure in current-clamp (n = 10) and voltage-clamp experiments (*n* = 7). Each neuron served as its own control, and *n* represents the number of recorded cells included in each comparison. Data are presented as mean ± standard error of the mean (SEM), unless otherwise indicated. Representative traces and figures were prepared using Clampfit 10.7 (Molecular Devices, USA) and GraphPad Prism 9 (GraphPad Software, USA). Statistical analyses were performed in Python using SciPy and statsmodels, and figures were generated using Matplotlib.

Treatment effects were quantified as the mean paired difference (*M*_diff_), calculated as the average within-neuron difference between treatment and control measurements (Treatment−Ctrl). Normality was assessed using the Shapiro–Wilk test applied to the paired differences. Paired comparisons were evaluated using two-tailed paired Students’s t-tests, with statistical significance defined as p < 0.05. To complement hypothesis testing, 95% confidence intervals (CIs) for *M*_diff_ were estimated using nonparametric paired bootstrap resampling with 10,000 iterations. In each iteration, neurons were sampled with replacement while preserving their paired control and treatment observations. Standardized effect sizes were displayed in forest plots, with corresponding 95% CIs obtained from 10,000 paired bootstrap resamples.

Data visualization combined paired individual-neuron plots with estimation plots displaying *M*_diff_, its 95% bootstrap CI, and the corresponding bootstrap sampling distribution. This estimation-based approach complements conventional hypothesis testing by emphasizing the magnitude and precision of treatment-associated changes (Ho et al., 2019; Martínez-Rojas et al., 2026).

## 3. Results

### H_2_S alters the passive membrane properties of CA1 pyramidal cells

To determine how exogenous H₂S influences the intrinsic electrical properties of CA1 PCs, we compared their electrophysiological responses under control conditions and following exposure to the H_2_S donor NaHS (200 µM). We began by characterizing their passive membrane responses in acute dorsal hippocampal slices using whole-cell patch-clamp recordings. Representative voltage traces (Fig. 1a) and the corresponding current–voltage (I–V) relationships (Fig. 1b) showed larger hyperpolarizing voltage responses to negative current injections after NaHS exposure. Consistent with this observation, NaHS exposure increased input resistance (*M*_diff_, 19.3 MΩ; 95% CI, 6.3–37.5 MΩ; p=0.049; Fig. 1c), as determined from the slope of each neuron’s I–V relationship. NaHS exposure also increased the membrane time constant (*M*_diff_, 14.1 ms; 95% CI, 8.5–21.0 ms; p=0.0025; Fig. 1d), estimated membrane capacitance (*M*_diff_, 83.6 pF; 95% CI, 38.7–120.5 pF; p=0.0045; Fig. 1e), and sag ratio (*M*_diff_, 0.09; 95% CI, 0.06–0.13; p=0.0010; Fig. 1f). In contrast, no statistically significant change in rheobase was detected (Fig. 1g; *see Supplementary Table 1 for detailed results).* Standardized effects sizes summarizes these changes in Fig. 1h. Together, these findings indicate that NaHS exposure increases the magnitude and prolongs the time course of membrane responses to hyperpolarizing current injections.

**Figure 1.**
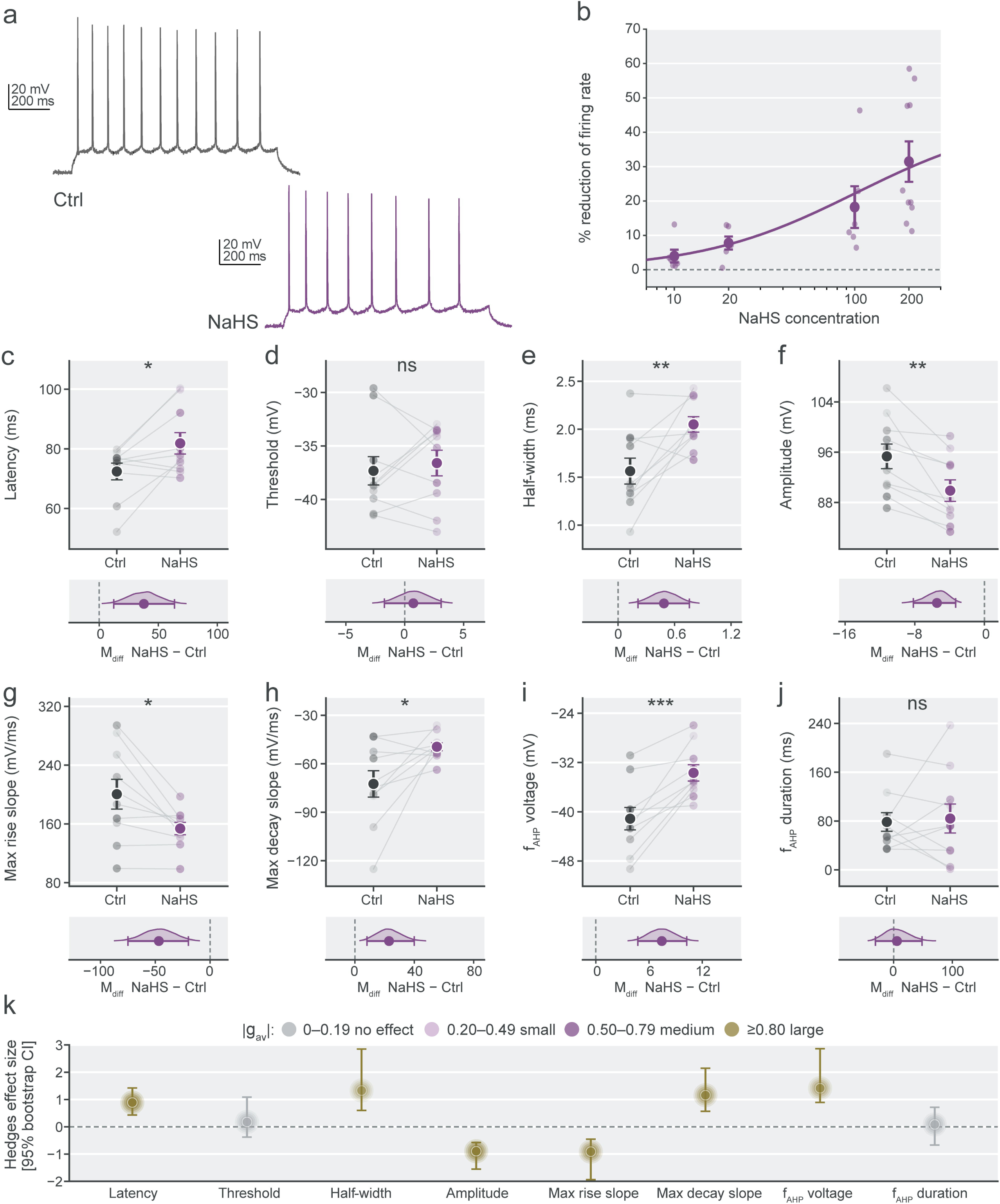
NaSH modifies passive membrane properties in hippocampal CA1 PCs. (a) Representative voltage responses to 1-s current steps ranging from −300 to +90 pA under control condition and from −300 to +150 pA after 5 min of NaHS perfusion (200 µM). (b) Current–voltage (I–V) relationships showing the effects of NaHS on membrane responses to current injections from 0 to −300 pA. (c–e) Paired measurements of somatic input resistance (c), membrane time constant (d), and membrane capacitance (e), characterizing changes in membrane responsiveness and charging dynamics. (f–g) Sag ratio (f) and rheobase (g), capturing distinct aspects of the membrane response to hyperpolarizing and depolarizing current injections, respectively. (h) Forest plot showing standardized effect sizes (Hedges’ |g_av_|) and their 95% bootstrap confidence intervals, with colors indicating effect-size magnitude based on absolute values. For this and subsequent figures, unless otherwise indicated, small symbols represent individual neurons, connecting lines indicate paired measurements. Large symbols and error bars represent mean ± S.E.M. Lower panels show the mean paired difference (*M*_diff_; NaHs – Ctrl) and the bootstrap sampling distribution. Confidence intervals for panels c-h were estimated using 10,000 paired bootstrap resamples. N = 10 individual neurons. Paired comparisons were assessed using two-tailed paired Student’s t-test. Asterisks indicate statistical significance: *p < 0.05; **p < 0.01; ***p < 0.001; ns, not significant.

### H_2_S modifies AP waveform and dynamics

Having established that H_2_S modifies membrane properties in CA1 PCs, we next examined its influence on AP firing and waveform dynamics. Representative recordings obtained with identical 1-s depolarizing current steps of 150 pA showed fewer APs during NaHS application than under control conditions (Fig. 2a). To examine the concentration dependence of this inhibition, firing responses were quantified across depolarizing current steps from 30 to 300 pA, and the integrated firing response was determined for each NaHS concentration (10, 20, 100, and 200 µM). Among the concentrations tested, 200 µM produced the greatest mean reduction, approximately 30% relative to control (Fig. 2b), and was selected for subsequent electrophysiological characterization.

**Figure 2.**
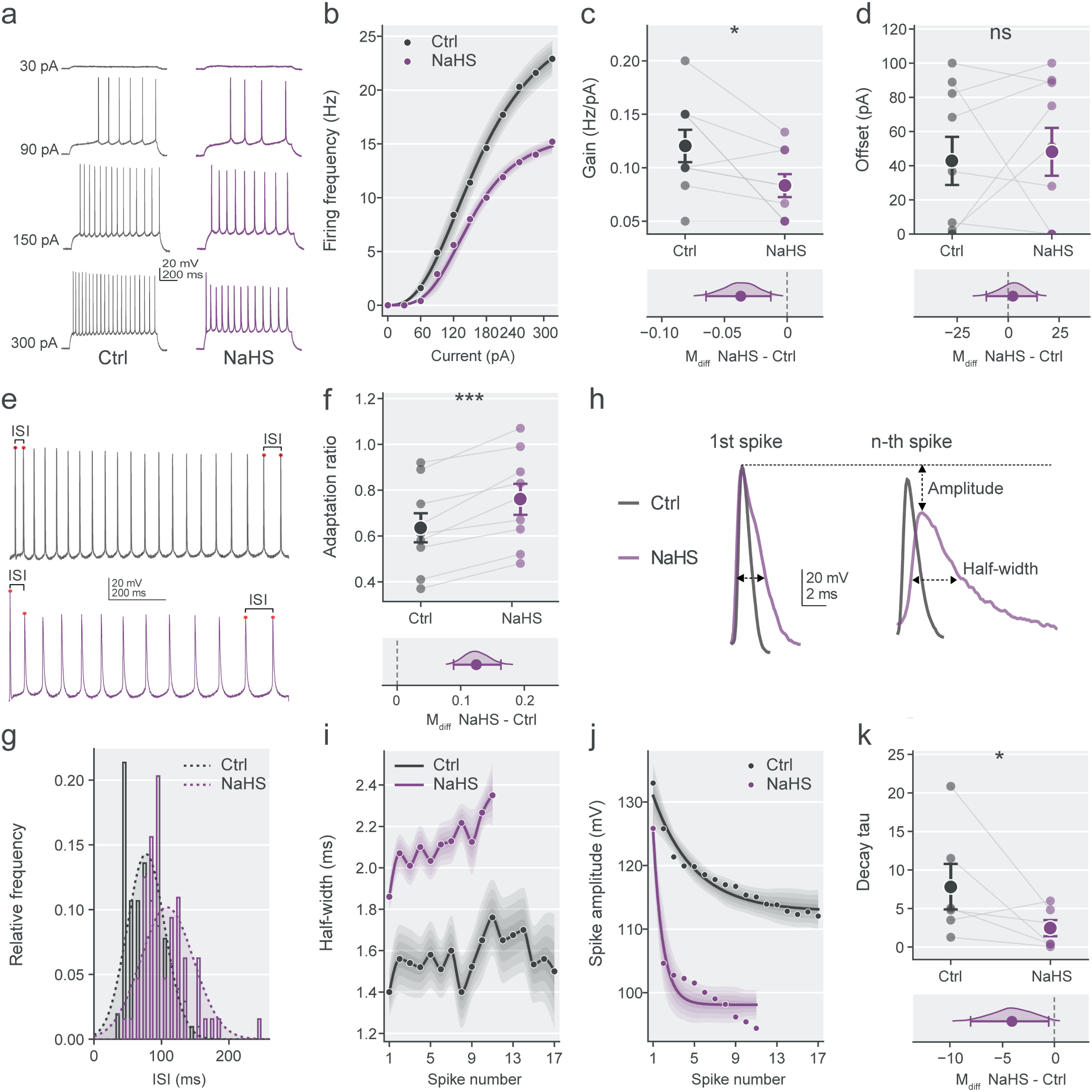
NaHS alters AP waveform dynamics of CA1 PCs. (a) Representative voltage responses to identical 1-s depolarizing current steps (150 pA) recorded before (Ctrl) and during NaHS application (200 µM). (b) Concentration–response relationship for NaHS (10, 20, 100, and 200 µM), quantified as the percentage reduction in the integrated firing response across depolarizing current injections from 30 to 300 pA, relative to control. (c–d) AP latency (c) and voltage threshold (Dd), characterizing spike initiation. (e–h) AP half-width (e), amplitude (f), maximum rise slope (g), and maximum decay slope (h), describing AP waveform and kinetics. (i–j) Fast afterhyperpolarization (f_AHP_) voltage (i) and duration (j). (k) Forest plot of standardized effect sizes (Hedges’ |g_av_|) with 95% bootstrap confidence intervals. Colors indicate effect-size magnitude based on absolute values. Measurements in panels c-j were obtained before and after 5 min of NaHS perfusion (200 µM). n = 10 neurons. Statistical conventions are as described in Figure 1.

At 200 µM, NaHS increased the latency to the first AP (*M*_diff_, 37.8 ms; 95% CI, 12.3–63.8 ms; p=0.0224; Fig. 2c), without statisticaly significant change in AP threshold (Fig. 2d). NaHS exposure also altered AP morphology in CA1 PCs, producing broader APs with reduced amplitude. Specifically, AP half-width increased (*M*_diff_, 0.5 ms; 95% CI, 0.2–0.8 ms; p=0.0088; Fig. 2e), whereas AP amplitude decreased (*M*_diff_, −5.5 mV; 95% CI, −8.2 to −3.3 mV; p=0.0027; Fig. 2f). Additionally, NaHS slowed AP kinetics by reducing the maximal rise slope (*M*_diff_, −46.8 mV/ms; 95% CI, −74.9 to −19.8 mV/ms; p=0.0119; Fig. 2g) and making the maximal decay slope less negative (−72.4 vs. −49.5 mV/ms; *M*_diff_, 23.0 mV/ms; 95% CI, 8.1–39.9 mV/ms; p=0.0245; Fig. 2h). The voltage at the fast afterhyperpolarization (f_AHP_) was likewise less negative (−41.1 vs. −33.7 mV; *M*_diff_, 7.4 mV; 95% CI, 4.7–10.3 mV; p=0.0008; Fig. 2i), with no statistically significant change in its duration (Fig. 2j). Standardized effect sizes summarize these changes in Fig. 2k. Together, these results show that NaHS exposure delays AP initiation and produces broader, smaller spikes with slower rising and falling phases.

### H_2_S attenuates neuronal input–output gain and alters AP properties during repetitive firing

To determine how H_2_S affects sustained firing across different levels of depolarizing input, current steps of increasing amplitude were applied to CA1 PCs. Representative current-clamp recordings at 30, 90, 150, and 300 pA illustrate the responses obtained under control conditions and during NaHS exposure (200 µM; Fig. 3a). Analysis of the neuronal input-output (I-O) relationship revealed a reduction in firing response during NaHS exposure (*M*_diff_, −0.03 Hz/pA; 95% CI, −0.06 to −0.01 Hz/pA; p=0.0304; Fig. 3c), with no statistically significant change in the offset (Fig. 3d). We next examined the temporal organization of AP trains. Expanded views of the 300-pA recordings revealed longer ISIs, during NaHS exposure (Fig. 3e). Consistent with these findings, the adaptation ratio increased (*M*_diff_, 0.12; 95% CI, 0.09–0.16; p=0.0003; Fig. 3f), and the ISI distributions shifted toward larger intervals (Fig. 3g). We also examined changes in AP amplitude and half-width throughout the AP train, as illustrated by comparisons of the first and later AP (Fig. 3h). Plotting these parameters against AP number revealed a progressive AP broadening during NaHS exposure (Fig. 3i) and a more rapid decline in amplitude across successive spikes (Fig. 3j). This decline was fitted with an exponential function, yielding a smaller decay constant (τ) during NaHS exposure (*M*_diff_, −5.4 ms; 95% CI, −10.6 to −0.7 ms; p=0.0415; Fig. 3k). Together, these findings show that NaHS exposure reduces the output and gain of CA1 PCs and alters AP waveform kinetics during repetitive firing.

**Figure 3.**
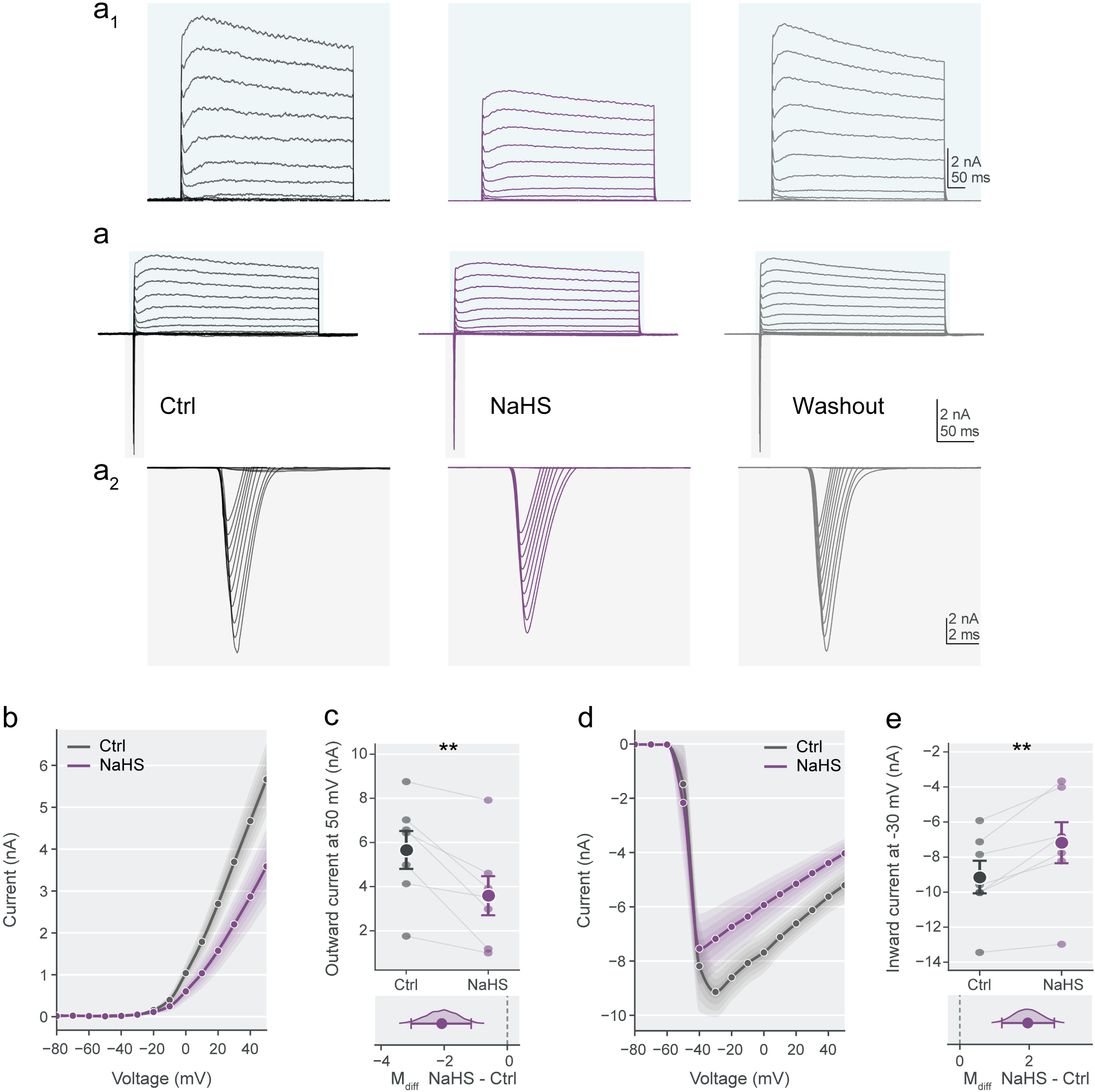
NaSH reduces firing frequency and neuronal input–output gain of CA1 PCs. (a) Representative current-clamp recordings from CA1 PCs under control (Ctrl) conditions and after 5 min of NaHS exposure (200 µM), showing AP trains evoked by 1-s depolarizing current injections of 30, 90, 150, and 300 pA. (b–d) Neuronal input–output (I–O) relationship showing firing frequency as a function of injected current (b), with paired comparisons of firing gain (c) and offset (d). (e–g) Magnified segments of the 300-pA recordings showing interspike intervals (ISIs) indicated on the traces (e), paired measurements of the adaptation ratio (f), and ISI distributions (g). (h–j) Comparison of the first and last APs within representative Ctrl and NaHS trains (h), with AP half-width (i) and amplitude (j) plotted as functions of spike number during 300-pA current injections. Solid curves in (j) represent exponential fits to the amplitude data. (k) Paired comparison of the exponential decay time constant (τ), expressed as the number of spikes and derived from the amplitude decay analysis in (j). n = 10 neurons. Statistical conventions are as described in Figure 1

### H_2_S suppresses voltage-dependent outward and inward currents of CA1 PCs

The H_2_S-induced slowing of AP depolarization and repolarization, together with the attenuation of sustained firing, suggested that voltage-dependent ionic currents underlying AP generation and repolarization may be affected. We therefore examined inward and outward membrane currents to investigate the current changes associated with these effects. We characterized voltage-dependent inward and outward currents under voltage-clamp configuration before and 5 min after NaHS exposure (200 µM), with additional recordings following washout to assess recovery (Fig. 4a). The outward and inward components are shown separately to resolve their respective responses to membrane depolarization (Fig. 4a_1_ and a_2_). The outward current-voltage (I-V) relationship showed that NaHS exposure reduced outward current amplitude at depolarized test potentials (Fig. 4b). To quantify this effect, we compared steady-state outward currents at +50 mV, which were significantly reduced during NaHS exposure (*M*_diff_, −2.1 nA; 95% CI, −3.1 to −1.2 nA; p=0.0078; Fig. 4c). Likewise, the inward I-V relationship showed reduced inward current magnitudes during NaHS exposure (Fig. 4d). Quantification of peak inward current at −30 mV confirmed that current became significantly less negative (*M*_diff_, 1.9 nA; 95% CI, 1.2 to 2.7 nA; p=0.0034; Fig. 4e). Together, these findings indicate that NaHS exposure reduces the magnitude of both inward and outward voltage-dependent membrane currents, providing an ionic correlate of the altered AP kinetics and reduced firing frequency observed under current clamp.

**Figure 4.**
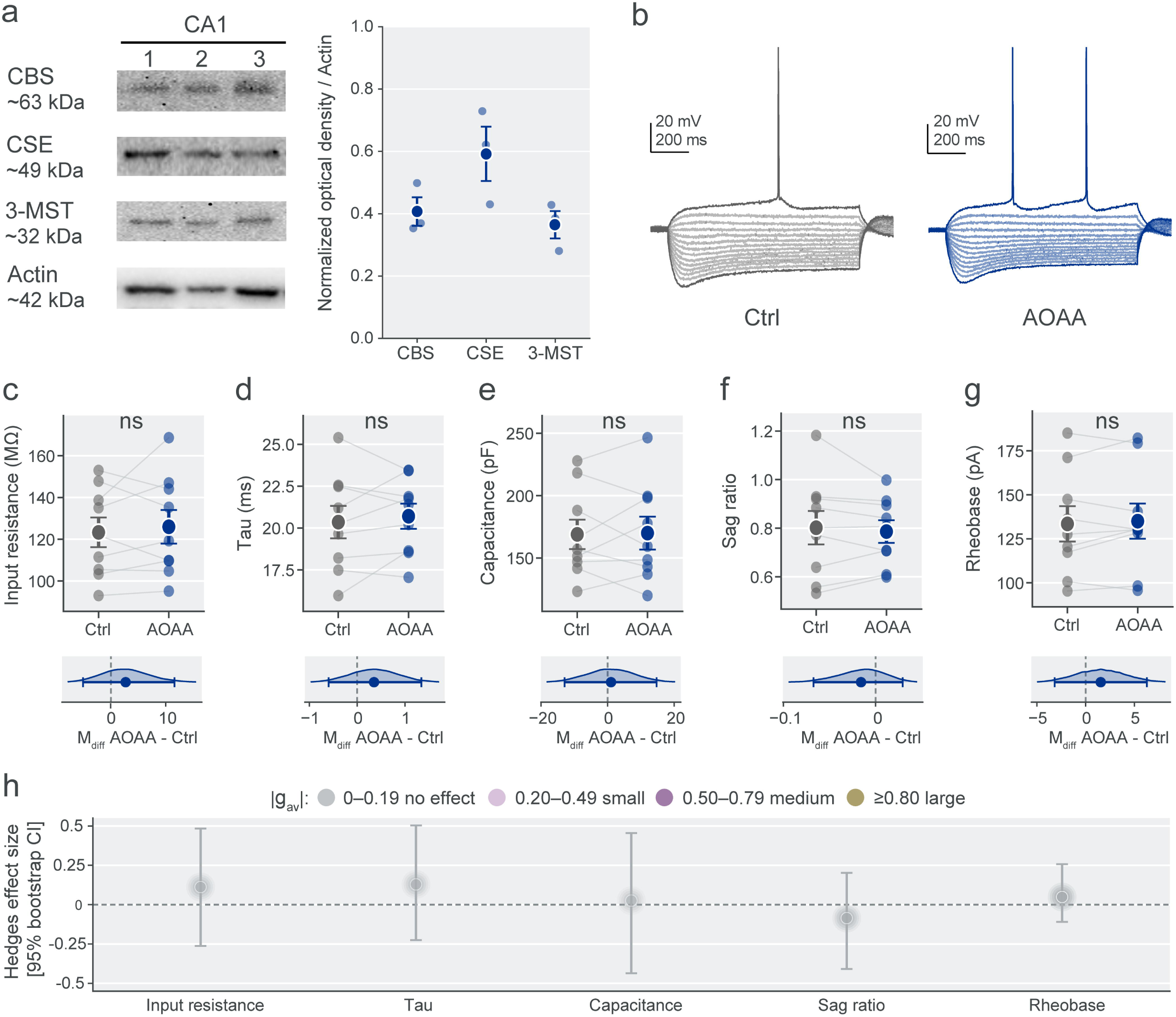
NaSH decreases voltage-dependent inward and outward currents in CA1 PCs. (a) Representative voltage-clamp recordings showing inward and outward membrane currents under control (Ctrl) conditions, after 5 min of NaHS exposure (200 µM), and following washout. Currents were elicited by voltage steps from −80 to +50 mV in 10 mV increments, from a holding potential of −65 mV. (a_1_, a_2_) Separate displays of the outward (a_1_) and inward (a_2_) current components from the recordings in (a). (b) Current–voltage (I–V) relationships of steady-state outward currents under control (Ctrl) and NaHS conditions across the voltage range examined. (c) Paired comparison of steady-state outward current amplitude at +50 mV. (d) I–V relationships of peak inward currents under Ctrl and NaHS conditions. (e) Paired comparison of peak inward current amplitude at −30 mV. N = 7 neurons. Statistical conventions are described in Figure 1.

### AOAA does not substantially alter passive membrane properties of CA1 pyramidal cells

Given the effects of exogenous H_2_S on AP dynamics, sustained firing, and voltage-dependent membrane currents, we next examined whether endogenous H_2_S synthesis contributes to the intrinsic electrical properties of CA1 PCs. To determine whether H_2_S enzymes were present in the stratum pyramidale, the anatomical region examined electrophysiologically, we assessed the expression of CBS, CSE, and 3-MST by Western blotting. Protein samples were obtained from microdissected tissue enriched in the CA1 stratum pyramidale rather than whole-hippocampal homogenates. Immunoreactive bands for all three enzymes were detected, and their signals were quantified relative to β-actin in samples from three independent naïve rats (Fig. 5a). We next examined membrane responses before and 5 min after perfusion with aminooxyacetic acid (AOAA, 100 µM), an inhibitor of pyridoxal phosphate-dependent enzymes that can interfere with endogenous H₂S production. Representative current-clamp recordings obtained with hyperpolarizing and depolarizing current steps are shown in Fig. 5b. No statistically significant changes were detected in somatic input resistance, membrane time constant, or membrane capacitance in CA1 PCs (Fig. 5c-e). Similarly, neither the sag ratio nor rheobase differed significantly between conditions (Fig. 5f-g). Standardized effect-size analysis further supported the consistency of this pattern across membrane properties: all five endpoints had Hedges’ *g*_av_ values near zero, and none met the predefined threshold for a small effect (|*g*_av_| = 0.20; Fig. 5h). Thus, although H₂S-related biosynthetic enzymes were detected in tissue enriched in the CA1 pyramidal layer, AOAA exposure did not produce detectable changes in the passive membrane properties or rheobase under the conditions examined.

**Figure 5.**
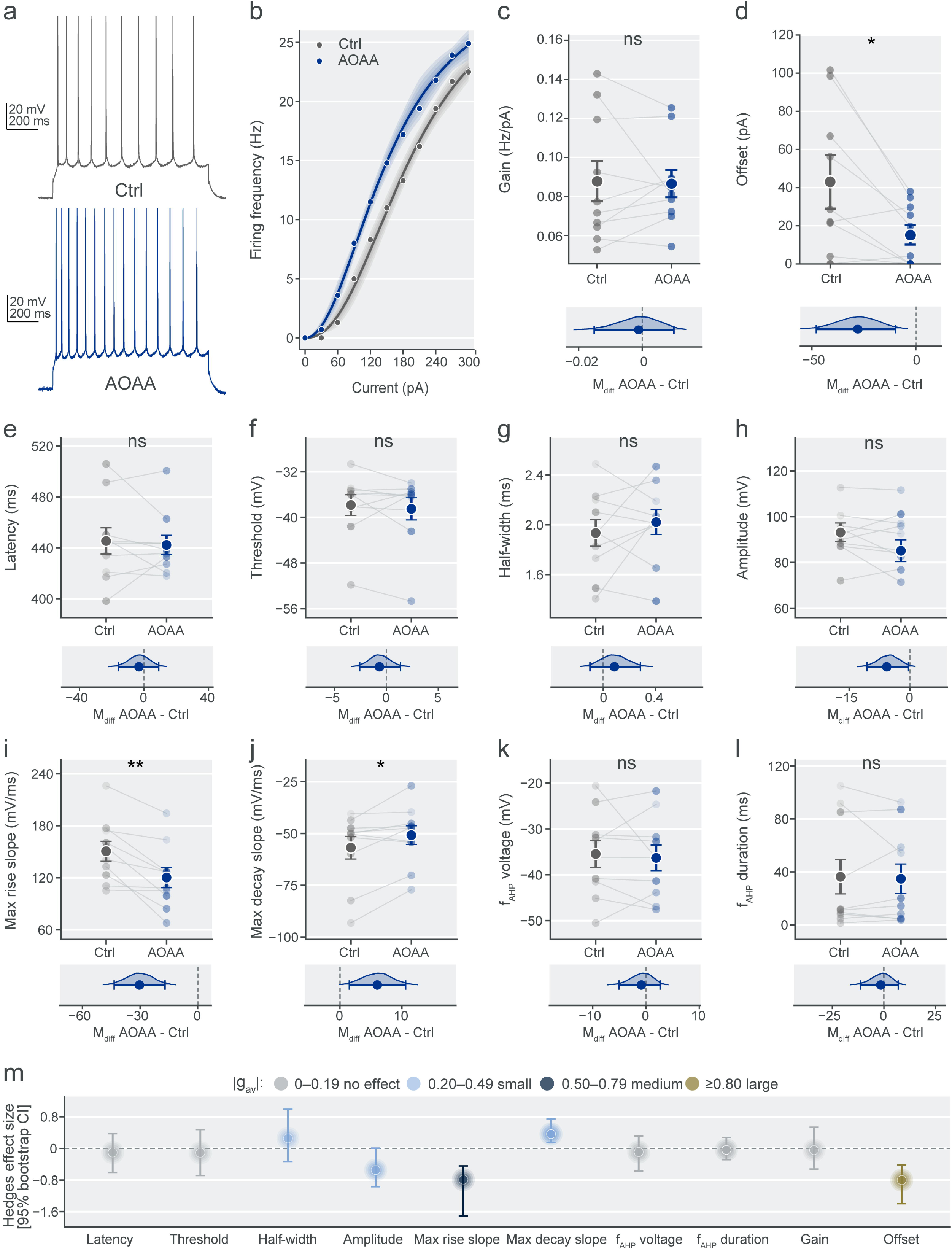
Expression of H_2_S-producing enzymes in the CA1 pyramidal layer and effects of AOAA on intrinsic membrane properties. (a) Representative Western blots showing cystathionine β-synthase (CBS), cystathionine γ-lyase (CSE), 3-mercaptopyruvate sulfurtransferase (3-MST), and βactin in tissue enriched in the CA1 stratum pyramidale, microdissected from three independent naïve rats. Corresponding densitometric quantification of each enzyme’s immunoreactive signal normalized to β-actin. Lanes 1-3 and individual symbols represent samples from the three animals, large symbols and error bars indicate mean ± S.E.M. (b) Representative current-clamp recordings from CA1 PCs under control (Ctrl) conditions and after 5 min of exposure to aminooxyacetic acid (AOAA, 100 µM), obtained with 1-s current steps ranging from −300 pA through progressively less hyperpolarizing injections to depolarizing pulses sufficient to evoke APs. (c–g) Paired comparisons of input resistance (c), membrane time constant (d), membrane capacitance (e), sag ratio (f), and rheobase (g) under Ctrl and AOAA conditions. (h) Forest plot summarizing standardized effect sizes (Hedges’ |g_av_|) and their 95% bootstrap confidence intervals for the five electrophysiological parameters. Colors within the symbols indicate effect-size magnitude based on absolute values. Statistical conventions are described in Figure 1.

### AOAA enhances firing in CA1 pyramidal cells

The lack of statistically detectable changes in passive membrane properties (Fig. 5) did not exclude changes in active excitability. We therefore examined the I-O relationship and AP kinetics under control conditions and during AOAA exposure (100 µM; Fig. 6a). AOAA increased the firing frequency of CA1 PCs, as reflected by an upward shift in I-O curve (Fig. 6b). Although no statistically significant change in neuronal gain was detected (Fig. 6c), AOAA significantly decreased the offset (*M*_diff_, −27.8 pA; 95% CI, −47.4 to −9.7 pA; p=0.0232; Fig. 6d), indicating a lower extrapolated current threshold for repetitive firing. Despite this shift, AP latency, threshold, half-width, and amplitude did not differ significantly from control during AOAA exposure (Fig. 6e–h). In contrast, AOAA reduced the maximum rise slope (*M*_diff_, −30.3 mV/ms; 95% CI, −43.5 to −17.0 mV/ms; p=0.0024; Fig. 6i) and the maximum decay slope (*M*_diff_, 6.0 mV/ms; 95% CI, 1.5 to 10.6 mV/ms; p=0.0351; Fig. 6j), indicating slower AP depolarization and repolarization. No statistically significant changes were detected in f_AHP_ voltage or duration (Fig. k-l). Standardized effect-size analysis summarize the changes in firing parameters and AP kinetics (Fig. 6m). Together, these findings demonstrate that AOAA increases firing frequency in CA1 PCs while slowing the rising and falling phases of APs.

**Figure 6.**
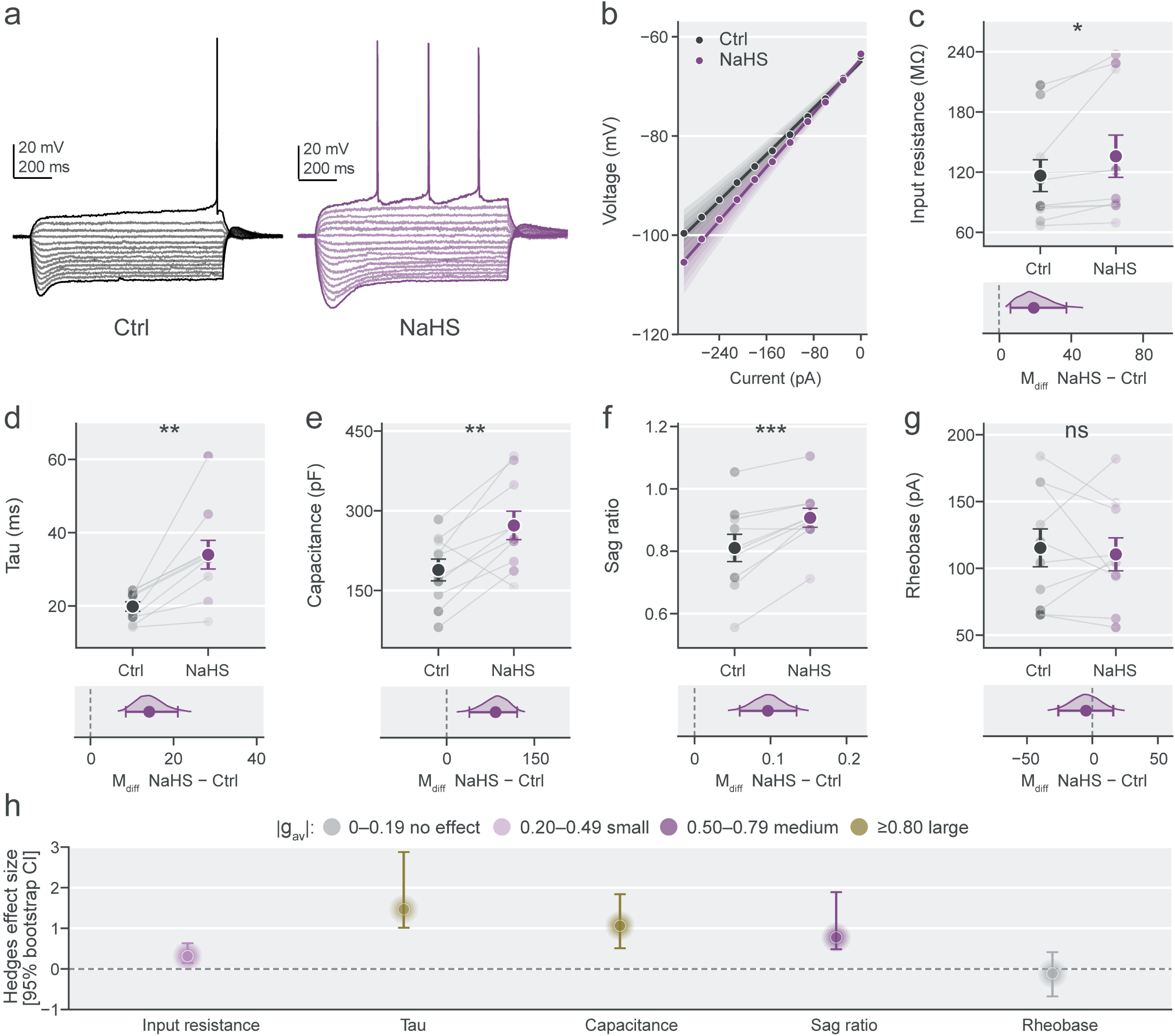
AOAA enhances firing frequency of CA1 PCs. (a) Representative current-clamp recordings showing AP firing under control (Ctrl) conditions and 5 min after AOAA exposure (100 µM). (b) Input–output (I–O) relationships showing firing frequency as a function of injected current under Ctrl and AOAA conditions. (c, d) Paired comparisons of firing gain (c) and offset (d), with corresponding mean-difference estimation plots. (e–l) Paired comparisons of AP latency (e), threshold (f), half-width (g), amplitude (h), maximum rise slope (i), maximum decay slope (j), fast afterhyperpolarization (f_AHP_) voltage (k), and f_AHP_ duration (l). (m) Forest plot summarizing standardized effect sizes (Hedges |g_av_|) and their 95% bootstrap confidence intervals for the eight AP parameters and for neuronal gain and offset. Forest plot colors indicate effect-size magnitude according to the absolute value of Hedges |g_av_|. Statistical conventions are described in Figure 1.

## 4. Discussion

The present study describes H₂S as a modulator of intrinsic excitability in CA1 PCs through two complementary pharmacological approaches: exogenous H₂S delivery using NaHS and pharmacological interference with endogenous H₂S biosynthesis using AOAA. H₂S altered intrinsic membrane properties, reduced firing output, and attenuated voltage-dependent inward and outward currents. Conversely, AOAA enhanced firing output without detectable changes in passive membrane properties. Together with the detection of H₂S-producing enzymes in the CA1 pyramidal layer, these findings support a functional relationship between sulfur-related pathways and the intrinsic mechanisms modulating hippocampal CA1 neuronal activity.

### H_2_S suppresses intrinsic excitability and alters the mechanisms supporting firing output

The electrophysiological changes induced by H_2_S converged on a reduced ability of CA1 PCs to sustain repetitive AP firing. At subthreshold potentials, the increase in somatic input resistance indicates a reduction in net membrane conductance. Previous studies have reported that NaHS increases (Khademullah and Ferguson, 2013) and decreases (Malik and Ferguson, 2016) the input resistance of distinct neuronal populations. In this study, we documented that H_2_S increased somatic input resistance, without enhancing neuronal output. Instead, the reduced I-O gain, together with the increased adaptation ratio and altered AP kinetics, suggests that H_2_S modulates the mechanisms that maintain sustained output of CA1 PCs. Thus, the increased input resistance does not compensate for the impaired firing response, which may be indicative of alterations in the responses mediated by voltage-dependent conductances involved in AP repolarization and recovery from inactivation. In CA1 PCs, BK-mediated repolarization supports high-frequency firing by limiting sodium-channel inactivation (Gu et al., 2007), whereas sodium-current availability critically regulates spike-frequency adaptation and neuronal gain (Fernandez et al., 2011). The involvement of voltage-gated currents in the effects of H_2_S on neuronal firing is consistent with previous studies reporting H_2_S-induced changes in neuronal excitability. In cultured hippocampal neurons, NaHS increased AP firing by inhibiting Kv2.1-mediated outward K⁺ currents (Dallas et al., 2021). Conversely, H_2_S reduced AP firing in sympathetic neurons through activation of K_ATP_ channels (Dominguez-Rodriguez et al., 2017). These contrasting findings highlight that the effects of H_2_S on neuronal firing depend on the specific ionic conductances modulated and their contribution to excitability. In this context, our voltage-clamp findings provided direct evidence of H_2_S-induced modulation of membrane currents associated with these electrophysiological changes. H_2_S significantly reduced, reversely, both inward and outward current amplitudes, consistent with the slower AP depolarization and repolarization slopes observed under current clamp.

On the other hand, representative recordings showed recovery of current amplitude following washout. These findings align with previous evidence that H_2_S modulates multiple classes of neuronal ion channels. In subfornical organ neurons, H_2_S enhanced transient and persistent voltage-gated Na^+^ currents and N-type Ca^2+^ currents, while leaving transient and delayed-rectifier K^+^ currents largely unaffected (Kuksis and Ferguson, 2015). In contrast, H_2_S inhibited sustained voltage-gated K^+^ currents in trigeminal ganglion neurons (Feng et al., 2013) and Kv2.1 channels in both native and recombinant systems (Dallas et al., 2021). These findings establish that H_2_S modulates voltage-dependent channels, suggesting that its effects depend on the specific ion channels expressed by each neuronal population. At the molecular level, H_2_S-mediated inhibition of Kv2.1 involves S-sulfhydration of cysteine 73, providing evidence for residue-specific regulation of channel function (Dallas et al., 2021). The redox-sensitive thiol groups of cysteine residues provide potential sites for sulfur-dependent modifications. Likewise, H₂S also activates TRPA1 channels through two intracellular cysteine residues located in their N-terminal domain (Ogawa et al., 2012). Given the functional role of astrocytic TRPA1 in the hippocampus (Shigetomi et al., 2013), this mechanism may also be relevant to sulfur-dependent regulation of hippocampal circuits. Moreover, H₂S can enhance NMDA receptor-mediated responses in hippocampal neurons (Abe and Kimura, 1996; Yakovlev et al., 2017), extending its regulatory actions to ligand-gated channels. Beyond receptor modulation, H_2_S increases extracellular levels of glutamate, GABA, and D-serine in the hippocampus (Furuie et al., 2023), suggesting that changes in the neurochemical environment may also contribute to H_2_S-induced alterations in CA1 neuronal activity. Together, these findings place our electrophysiological observations within a broader framework of H_2_S-dependent regulation of ion channels and neurotransmitter availability. Nevertheless, the reduced inward and outward currents observed under voltage clamp point to changes in membrane conductances whose molecular identities require further pharmacological and biophysical characterization.

### Endogenous H_2_S-Producing Enzymes in CA1 and AOAA-Induced Changes in Neuronal Excitability

Our Western blot analysis performed in tissue enriched in the pyramidal cell layer confirmed the expression of the principal enzymes involved in H₂S biosynthesis (Cirino et al., 2023). The blots indicate that this tissue contains the molecular machinery required for H_2_S synthesis. It would be of interest to explore the role of H_2_S beyond PCs, for example in local interneurons or astrocytes, as a previous study demonstrated CBS expression in astrocytes (Enokido et al., 2005). Interestingly, no information regarding the action of H2S on GABAergic interneurons is available. Overall, the expression of these enzymes supports the notion that the CA1 region of the hippocampus has the capacity for local H₂S production, although protein abundance does not establish enzymatic activity or identify the cellular sources of H₂S within CA1.

Interestingly, pharmacological interference with endogenous H_2_S synthesis using AOAA, did not noticeably alter membrane properties and produced changes in AP rise and decay slopes waveforms but increased the firing frequency of CA1 PCs, indicating a net enhancement of excitability during sustained depolarization. The lack of detectable changes in passive membrane properties suggest that H_2_S does not act on ion channels opened near the resting membrane potential and suggest instead, that H_2_S acts on voltage-dependent conductances, although the specific currents involved remain unresolved. Changes in neurotransmitter release may also contribute, given previous evidence that H_2_S promotes glutamate, GABA and D-serine release (Furuie et al., 2023).

The AOAA effect contrasts with the suppression of firing frequency induced by H_2_S but does not necessarily reflect an opposing action on endogenous H_2_S signaling. Given the multiple enzymatic targets of AOAA, the increased firing should not be attributed exclusively to inhibition of endogenous H_2_S synthesis. In addition to inhibiting CBS and CSE, AOAA can indirectly suppress the 3-MST pathway by inhibiting cysteine aminotransferase. AOAA also inhibits other pyridoxal-5′-phosphate-dependent enzymes, including GABA transaminase, glutamate decarboxylase, and aspartate aminotransferase (Zuhra et al., 2020). Thus, the AOAA-induced increase in neuronal firing may reflect reduced H_2_S synthesis, alterations in neurotransmitter or cellular metabolism, or a combination of these effects.

## Limitations

The uncertainty in the effective H_2_S concentration reaching CA1 PCs during NaHS perfusion is a central limitation of this study. The nominal donor concentration does not directly indicate the local availability of free sulfide, which exists predominantly as H_2_S and HS⁻ in a pH-dependent equilibrium. At pH 7.4 and 37 °C, approximately 20% of dissolved free sulfide is present as molecular H₂S, while its availability is further influenced by volatilization, oxidation, and tissue metabolism (Olson, 2012). Although continuous perfusion supplies H_2_S to the slice, diffusion into the tissue and local consumption may establish concentration gradients between the bathing solution and recorded neurons. Consequently, neither the extracellular concentration at the recording site nor the intracellular H_2_S exposure can be inferred directly from the nominal NaHS concentration. Estimates of endogenous brain H_2_S concentrations vary substantially with the analytical method employed. Earlier studies reported tissue H_2_S concentrations of 47–166 μM, whereas subsequent measurements placed free H_2_S levels in mouse brain at approximately 15 nM (Furne et al., 2008). This discrepancy highlights the difficulty of defining physiological sulfide exposure and underscores the importance of distinguishing nominal NaHS concentrations from the H_2_S available to CA1 neurons during perfusion. Beyond its physiological actions, sulfide at higher exposures may compromise neuronal bioenergetics through inhibition of mitochondrial cytochrome c oxidase, a well-established mechanism of sulfide toxicity (Nicholson et al., 1998). Although metabolic impairment could contribute to the reduced firing output observed after H₂S exposure, mitochondrial function was not assessed in the present study. Thus, the observed electrophysiological changes should not, in themselves, be interpreted as evidence of neurotoxicity.

## Conclusion

Our findings establish intrinsic membrane excitability as an important target of H₂S modulation in hippocampal CA1 PCs. By influencing the voltage-dependent processes that determine AP generation and sustained firing, H₂S can alter not only the electrical properties of individual APs but, more fundamentally, the relationship between depolarizing input and neuronal output. This integrated effect highlights how changes in membrane conductances can reshape the capacity of CA1 neurons to sustain activity in response to excitatory drive. Although the specific ionic and metabolic contributions remain to be established, our results provide an electrophysiological basis for understanding how H₂S may influence neuronal responsiveness and, potentially, the functional behavior of hippocampal circuits.

## Acknowledgements

The authors acknowledge financial support from CINVESTAV, Mexico.

## Data statement

The data supporting the findings of this study are available from the corresponding author upon reasonable request.

## CRediT authorship contribution statement

Martínez-Rojas VA: Conceptualization, Methodology, Investigation, Formal analysis, Writing, review, and editing; Perez-Barragan A: Methodology, Investigation, Formal analysis and Writing; Rojas-Hernández L: Formal analysis; Ramírez-Camacho I: Methodology, Investigation; Centurion D: Conceptualization, Investigation; Galván EJ: Conceptualization, Methodology, Investigation, Formal analysis, Writing, review, and editing.

