## Supplementary Table 1 for "Hydrogen sulfide modulation of intrinsic excitability in rat hippocampal CA1 pyramidal cells"

| **Figure** | | **Variable** | **Control mean ± SEM** | **Treatment mean ± SEM** | **Δ mean** | **95% CI of Δ** | **p** | **Hedges gₐᵥ** | **95% CI of g** |
| --- | --- | --- | --- | --- | --- | --- | --- | --- | --- |
| 1 | c | Input resistance (MΩ) | 116.504 ± 15.814 | 135.817 ±  21.077 | 19.31 | [6.314, 37.519] | 0.0493 | 0.314 | [0.146, 0.641] |
| 1 | d | Tau (ms) | 19.833 ±  1.283 | 33.977 ±  3.921 | 14.14 | [8.505, 21.038] | 0.0025 | 1.468 | [1.018, 2.944] |
| 1 | e | Capacitance (pF) | 188.731 ± 20.451 | 272.337 ±  26.822 | 83.60 | [38.712, 120.527] | 0.0045 | 1.062 | [0.512, 1.853] |
| 1 | f | Sag ratio | 0.811 ±  0.044 | 0.907 ±  0.031 | 0.09 | [0.059, 0.134] | 0.0010 | 0.776 | [0.482, 1.865] |
| 1 | g | Rheobase (pA) | 115.296 ± 14.206 | 110.434 ±  12.308 | -4.86 | [-25.677, 15.945] | 0.6780 | -0.111 | [-0.663, 0.408] |
| 2 | c | Latency (ms) | 209.585 ± 11.258 | 247.376 ±  14.318 | 37.79 | [12.335, 63.771] | 0.0224 | 0.889 | [0.431, 1.424] |
| 2 | d | Threshold (mV) | -37.329 ±  1.324 | -36.599 ±  1.200 | 0.73 | [-1.721, 3.091] | 0.5869 | 0.175 | [-0.379, 1.086] |
| 2 | e | Half-width (ms) | 1.563 ±  0.135 | 2.051 ±  0.081 | 0.48 | [0.212, 0.758] | 0.0088 | 1.329 | [0.598, 2.848] |
| 2 | f | Amplitude (mV) | 95.328 ±  1.960 | 89.877 ±  1.714 | -5.45 | [-8.158, -3.294] | 0.0027 | -0.897 | [-1.554, -0.576] |
| 2 | g | Max rise slope  (mV/ms) | 200.466 ± 20.312 | 153.694 ±  8.370 | -46.77 | [-74.945, -19.782] | 0.0119 | -0.912 | [-1.940, -0.456] |
| 2 | h | Max decay slope (mV/ms) | -72.440 ±  8.131 | -49.452 ±  2.501 | 22.98 | [8.082, 39.931] | 0.0245 | 1.157 | [0.566, 2.147] |
| 2 | i | f_AHP_ voltage (mV) | -41.127 ±  1.822 | -33.686 ±  1.331 | 7.44 | [4.734, 10.245] | 0.0008 | 1.412 | [0.891, 2.861] |
| 2 | j | f_AHP_ duration (ms) | 78.493 ±  15.259 | 84.111 ±  23.857 | 5.61 | [-30.957, 48.338] | 0.7997 | 0.085 | [-0.672, 0.713] |
| 3 | c | Gain (Hz/pA) | 0.120 ±  0.015 | 0.083 ±  0.011 | -0.03 | [-0.065, -0.013] | 0.0304 | -0.895 | [-1.875, -0.352] |
| 3 | d | Offset (pA) | 42.778 ±  14.071 | 48.111 ±  14.027 | 5.33 | [-24.034, 31.984] | 0.7354 | 0.121 | [-0.576, 0.889] |
| 3 | f | Adaptation ratio | 0.636 ±  0.063 | 0.760 ±  0.068 | 0.12 | [0.090, 0.162] | 0.0003 | 0.603 | [0.420, 1.140]] |
| 3 | k | Spike-amplitude decay | 7.830 ±  2.955 | 2.458 ±  1.045 | -5.37 | [-10.601, -0.712] | 0.0415 | -0.913 | [-2.074, -0.269 |
| 4 | c | Outward current (nA) | 5.665 ±  0.858 | 3.588 ±  0.883 | -2.07 | [-3.069, -1.155] | 0.0078 | -0.844 | [-2.523, -0.397] |
| 4 | e | Inward current  (nA) | -9.140 ±  0.930 | -7.183 ±  1.171 | 1.95 | [1.216, 2.698] | 0.0034 | 0.655 | [0.347, 1.777] |
| 5 | A | CBS / actin (normalized OD) | — | 0.407 ±  0.046 | — | — | — | — | — |
| 5 | A | 3-MST / actin (normalized OD) | — | 0.365 ±  0.044 | — | — | — | — | — |
| 5 | A | CSE / actin (normalized OD) | — | 0.592 ±  0.087 | — | — | — | — | — |
| 5 | c | Input resistance (MΩ) | 123.284 ±  7.100 | 125.959 ±  8.031 | 2.67 | [-5.105, 11.583] | 0.5689 | 0.112 | [-0.262, 0.484] |
| 5 | d | Tau (ms) | 20.352 ±  0.972 | 20.704 ±  0.750 | 0.35 | [-0.595, 1.342] | 0.5180 | 0.129 | [-0.226, 0.585] |
| 5 | e | Capacitance (pF) | 168.932 ± 11.808 | 169.956 ±  13.156 | 1.02 | [-12.924, 14.732] | 0.8955 | 0.026 | [-0.436, 0.455] |
| 5 | f | Sag ratio | 0.802 ±  0.069 | 0.786 ±  0.046 | -0.01 | [-0.067, 0.029] | 0.5680 | -0.085 | [-0.409, 0.203] |
| 5 | g | Rheobase (pA) | 133.480 ± 10.054 | 135.025 ±  10.024 | 1.54 | [-3.184, 6.286] | 0.5655 | 0.049 | [-0.109, 0.257] |
| 6 | c | Gain (Hz/pA) | 0.088 ±  0.010 | 0.087 ±  0.007 | -0.001 | [-0.015, 0.010] | 0.8574 | -0.043 | [-0.523, 0.565] |
| 6 | d | Offset (pA) | 43.02 ±  13.99 | 15.19 ±  5.04 | −27.83 | [−47.375, −9.715] | 0.0232 | -0.801 | [-1.372, -0.411] |
| 6 | e | Latency (ms) | 445.361 ± 10.320 | 442.239 ±  7.667 | -3.12 | [-15.721, 9.179] | 0.6523 | -0.104 | [-0.608, 0.374] |
| 6 | f | Threshold (mV) | -37.837 ±  1.792 | -38.510 ±  1.946 | -0.67 | [-2.585, 1.361] | 0.5388 | -0.109 | [-0.687, 0.476] |
| 6 | g | Half-width (ms) | 1.934 ±  0.106 | 2.020 ±  0.099 | 0.08 | [-0.100, 0.285] | 0.4314 | 0.254 | [-0.329, 0.989] |
| 6 | h | Amplitude (mV) | 93.125 ±  4.094 | 85.113 ±  4.729 | -8.01 | [-13.299, -3.116] | 0.0516 | -0.549 | [-1.248, -0.273] |
| 6 | i | Max rise slope (mV/ms) | 150.613 ± 11.516 | 120.297 ±  11.732 | -30.31 | [-43.460, -17.031] | 0.0024 | -0.790 | [-1.710, -0.444] |
| 6 | j | Max decay slope (mV/ms) | -56.834 ±  5.463 | -50.812 ±  4.544 | 6.02 | [1.518, 10.592] | 0.0351 | 0.363 | [0.151, 0.747] |
| 6 | k | f_AHP_ voltage (mV) | -35.464 ±  2.930 | -36.328 ±  2.798 | -0.86 | [-5.088, 2.668] | 0.6876 | -0.091 | [-0.576, 0.309] |
| 6 | l | f_AHP_ duration (ms) | 36.292 ±  12.987 | 34.783 ±  11.150 | -1.50 | [-11.450, 6.976] | 0.7683 | -0.038 | [-0.285, 0.279] |

Values are mean ± SEM; Δ = treatment − control. Confidence intervals are 95% paired-bootstrap intervals. p values are from two-sided paired t-tests. Figure 5 Western blot values are only descriptive.
